# Inhibition of both mutant and wild-type RAS-GTP in KRAS G12C colorectal cancer through cotreatment with G12C and EGFR inhibitors

**DOI:** 10.1101/845263

**Authors:** Thomas McFall, Michael Trogdon, Laura Sisk-Hackworth, Edward C. Stites

## Abstract

The combination of KRAS G12C inhibitors with EGFR inhibitors has reproducibly been shown to be beneficial. Here, we reveal a new benefit of this combination: it effectively inhibits both wild-type and mutant RAS. A role for WT RAS inhibition has not previously been reported for this important combination of targeted therapies. We believe that targeting both mutant and wild-type RAS helps explain why this combination of inhibitors is effective.

## INTRODUCTION

KRAS inhibitor, AMG-510 ^1^ (brand name Lumakras, generic name sotorasib) recently obtained FDA approval for the treatment non-small cell lung cancer patients who have the KRAS G12C mutation. Its utility on other forms of cancer with the G12C mutation is still being investigated. Colorectal cancer with the G12C mutation has appeared particularly intractable to G12C inhibition, motivating a desire to identify combination strategies and their mechanism of action ^2,3^.

Many different studies have identified the combination of EGFR inhibitors and KRAS G12C inhibitors as particularly effective in a variety of cancer types ^1–8^. That co-treatment with EGFR inhibitors and KRAS inhibitors would be effective was initially seen as counterintuitive ^9,10^. The first mechanism proposed to explain why the combination is effective was that blocking EGFR reduces the GTP-loading of KRAS G12C by RAS GEFs. As the G12C inhibitor in that study was also shown to only be capable of inhibiting the GDP bound form of RAS, increased levels of GDP bound KRAS G12C would in turn promote better inhibition of KRAS G12C ^4,5^. Another reported mechanism involves counteracting the loss of negative feedback after KRAS G12C inhibition. Strong RAS pathway oncogenes drive strong signaling from ERK, which in turn results in the activation of negative feedback pathways. The treatment of the RAS pathway oncogene with a targeted therapy drops the ERK signal and causes a loss of negative feedback, which may reactivate parts of the pathway. EGFR inhibitors have also been proposed to have a role in blocking the reactivation of EGFR after the loss of RAS pathway negative feedback ^1–3,8^. Whether reduced GTP-loading on KRAS G12C and reduced EGFR reactivation are the only mechanisms to explain the benefit of combining an EGFR inhibitor with KRAS G12C inhibitor is unclear.

## RESULTS

We investigated the combination of KRAS G12C and EGFR inhibitors. We focus on the combination of AMG-510, the first KRAS G12C inhibitor to receive FDA approval, with cetuximab, the EGFR inhibitor that is regularly utilized for colorectal cancer patients ^11^. Our combination treatment drug dose response experiments detected low levels of proliferation for the combination of EGFR inhibitors with G12C inhibitors for all three of the KRAS G12C mutant cell lines (**Figure 1A**). Using the excess over bliss score as a measure of synergy ^12^, we found that there was synergy for all three KRAS G12C cell lines (**Figure 1B**). Overall, these experiments further support the idea that the combination of G12C and EGFR inhibitors may add value over G12C inhibitor alone.

**Figure 1.**
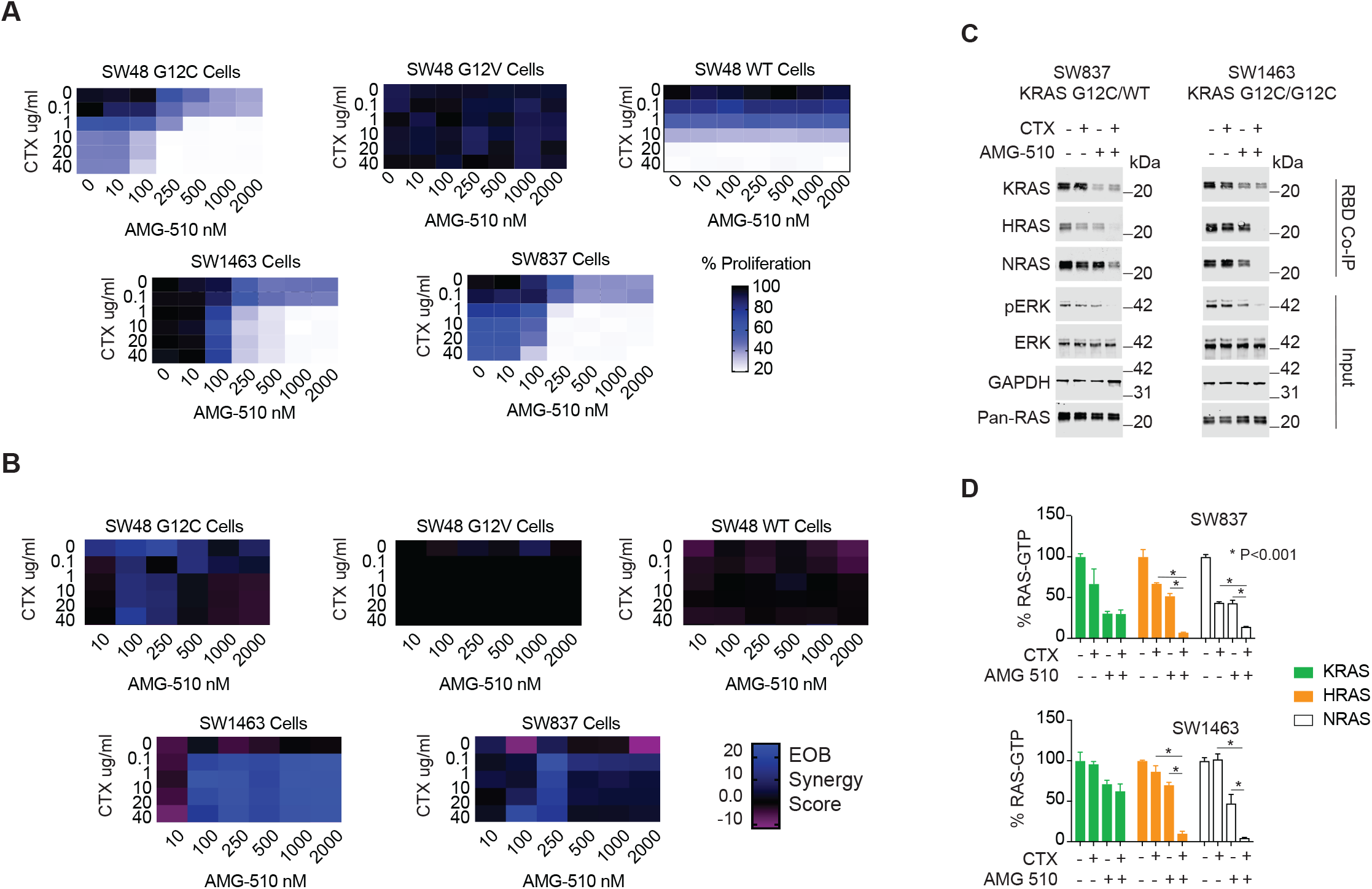
The combination of AMG-510 and cetuximab results in suppression of wild-type RAS-GTP. **(A**) Cellular proliferation as measured by the MTT assay for SW48 KRAS G12C^+/−^, SW48 KRAS G12V^+/−^, SW48 KRAS WT, SW1463 (KRAS G12C ^+/+^), and SW837 (KRAS G12C^+/−^) treated with combinations of cetuximab and AMG-510 for 48 hours. Heatmaps present average values from three separate experiments. (**B**) Calculated Excess over Bliss (EOB) synergy scores for the data in (A). (**C**) Active RAS measurements by RBD-WB for SW837 and SW1463 cells treated either with vehicle, 1μg/ml cetuximab, 250nM AMG-510 or both 1μg/ml cetuximab and 250nM AMG-510 for 48 hours. Results are representative of three separate experiments. (**D**) Mean RAS-GTP levels from three RBD-WB RBD pulldown assays with standard deviation. IP and western blot experiments were performed three times, bar plots represent mean abundance and error bars represent standard deviation. Statistical significance was determined by performing One-way ANOVA followed by post-hoc Tukey’s test for multiple comparisons, P-values are indicated within each panel comparing combination treatment to cetuximab and AMG-510 monotherapeies.

In our recent work we have demonstrated EGFR inhibitor sensitive KRAS mutants display reduced wild-type RAS-GTP upon treatment with EGFR inhibitors ^13–15^. To our knowledge, a role for WT RAS-GTP has not been evaluated for the combination of G12C and EGFR inhibitors. We therefore set out to experimentally evaluate how HRAS-, NRAS-, and KRAS-GTP levels change upon treatment with each inhibitor alone and in combination.

We observed profound suppression of KRAS-, NRAS-, and HRAS-GTP when the G12C mutant cells were treated with both cetuximab and AMG-510 (**Figure 1C,D**). That KRAS-G12C-GTP is more effectively suppressed by the combination of a G12C inhibitor with an EGFR inhibitor has been observed previously in multiple studies ^1–8^. However, we also found this combination to suppress WT RAS-GTP (i.e. HRAS-GTP and NRAS-GTP), and such a role has not previously been described. This suppression of WT RAS-GTP and KRAS-G12C-GTP was accompanied by an increased reduction in ERK phosphorylation (**Figure 1C**).

We wanted to further investigate the phenomenon of co-targeting WT and mutant RAS-GTP, which is a novel concept for the combination of G12C and EGFR inhibitors. We have a unique tool for studying oncogenic RAS and its targeting: a mathematical model of RAS signal regulation that relates biochemical defects of RAS mutants to observable cellular levels of RAS-GTP ^16^ (**Figure 2A**). We had previously updated this model to study the early G12C inhibitors ^17^, and we desired to use the model to study the combination of EGFR and G12C inhibitors. To do this, we needed to updated our KRAS G12C and KRAS G12C inhibition model for AMG-510.

**Figure 2.**
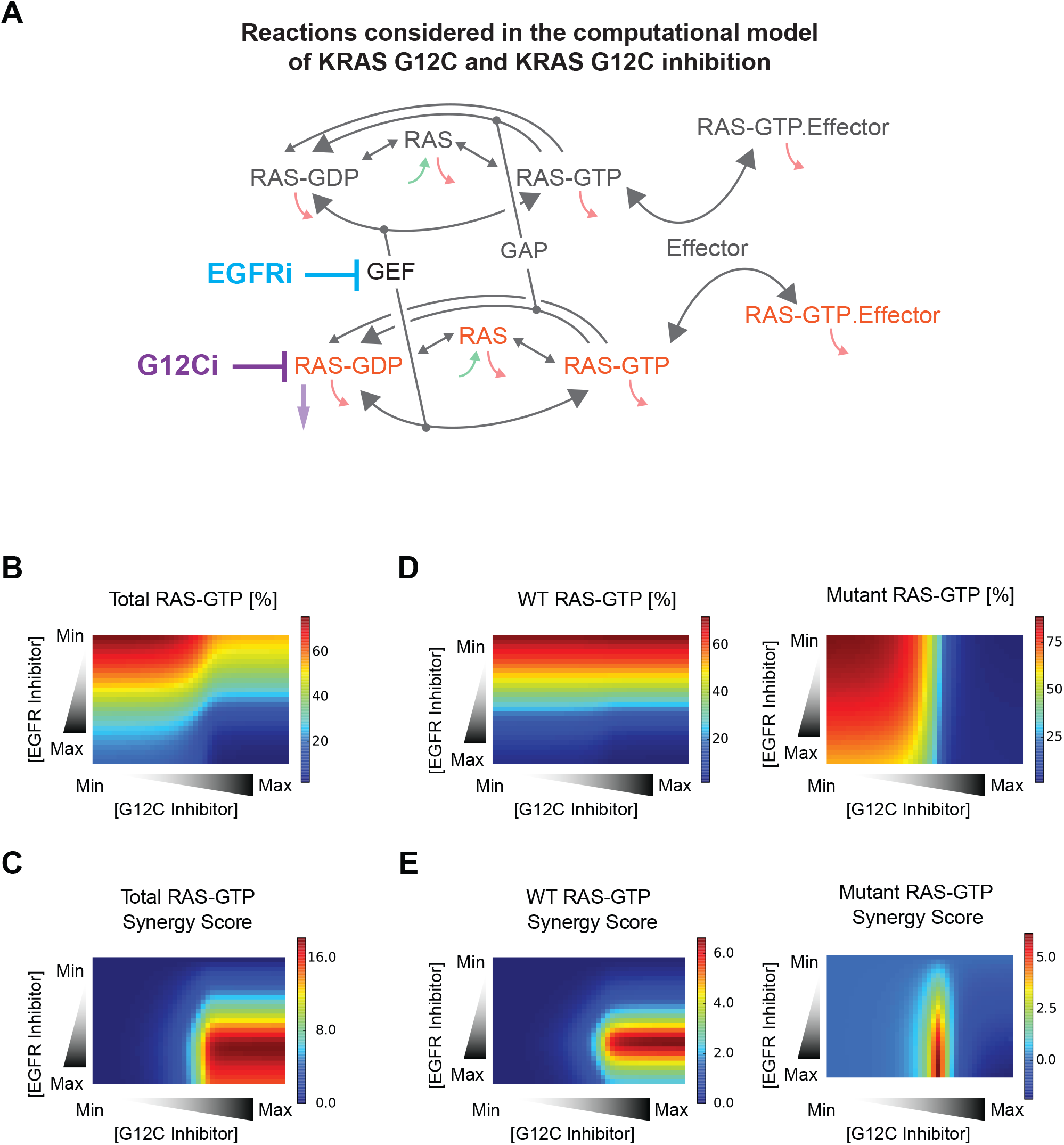
Mathematical modeling of KRAS G12C inhibition by the KRAS G12C and EGFR inhibitors finds the biophysical properties of the RAS network proteins and the RAS and EGFR inhibitors are sufficient to explain the co-targeting wild-type and mutant RAS. (**A**) Schematic of model of RAS regulation and KRAS G12C inhibition. Arrow colors indicate the type of reactions: gray, biochemical reactions that regulate the RAS nucleotide binding state; green, RAS production; red, RAS degradation. (**B**) Computational model-predicted levels of total RAS-GTP for the combination of KRAS G12C inhibition and EGFR inhibition. (**C**) Computational model-predicted Excess over Bliss (EOB) synergy scores based on predicted total RAS-GTP for the combination of KRAS G12C inhibition and EGFR inhibition. (**D**) Model-predicted levels of mutant RAS-GTP and WT RAS-GTP for the combination of KRAS G12C inhibition and EGFR inhibition for a modeled KRAS G12C heterozygous network. (**E**) Model-predicted EOB synergy scores based on predicted mutant RAS-GTP levels and WT RAS-GTP levels.

Previously described KRAS G12C inhibitors are believed to lock the KRAS G12C in the GDP-bound, inactive state and render it unable to interact with proteins that specifically bind to the GTP-bound conformation of RAS ^4,5^. We investigated whether AMG-510 also disrupts interactions between the KRAS-G12C protein with CRAF and to NF1, which we assessed a Bioluminescence Resonance Energy Transfer (BRET) assay. We observed KRAS-G12C treated with AMG-510 interacted much less with CRAF (**Figure S1A**), and that KRAS-G12C treated with AMG-510 interacted much less with NF1 (**Figure S1B**). We further confirmed that AMG-510 disrupts these interactions with a co-immunoprecipitation assay (**Figure S1C,D**). Thus, we concluded that the structure and equations of our model applied to AMG-510. We did need to update our parameters for the use of a different drug. AMG-510 has been described to engage KRAS G12C much more rapidly than the earlier KRAS G12C inhibitors; i.e. 9900/Ms k_inact_/K_i_ for AMG-510 ^18^ compared to 76/Ms for ARS-853 ^5^. We also updated our model to include a recently described reduction in the strength of the interaction between KRAS G12C and NF1 ^15^. Simulations with the updated model find AMG-510 more effective at inhibiting KRAS G12C than ARS-853 (**Figure S2**).

We then simulated combinations of EGFR and KRAS G12C inhibitors. The computational model reveals that the known mechanism of the inhibitors, the known properties of the drug, and the modeled key reactions of the RAS signaling network are sufficient to reproduce the added synergistic benefits of co-targeting EGFR and KRAS G12C (**Figure 2B,C**). Our simulations model the reduction in both WT and mutant RAS-GTP levels (**Figure 2D**), and our simulations find that there will also be considerable levels of synergy for both WT RAS-GTP and mutant RAS-GTP (**Figure 2E).**Overall, the model suggests that synergy in the combination of G12C inhibitors and EGFR inhibitors has contributions targeting both from mutant and WT RAS-GTP.

Previously demonstrated benefits of co-targeting KRAS G12C and EGFR, which involve increased loading of the G12C inhibitor and targeting EGFR after negative feedback pathways are released, are all also likely to be at play. Our work presents a third benefit of this combination: combined targeting of WT and mutant RAS. To the best of our knowledge, we are the only group to investigate the effects of G12C inhibitor and EGFR inhibitor combinations on WT HRAS and WT NRAS signaling, which we have shown here to be a critical component of the response to combination treatment in colorectal cancer cells. That wild-type RAS signaling is frequently elevated in the presence of mutant RAS has extensive experimental evidence ^13,16,19–21^, yet it is an aspect of RAS signaling that is often overlooked when mutant RAS cancers are considered ^22^.

---

**Supplementary Figure 1.**
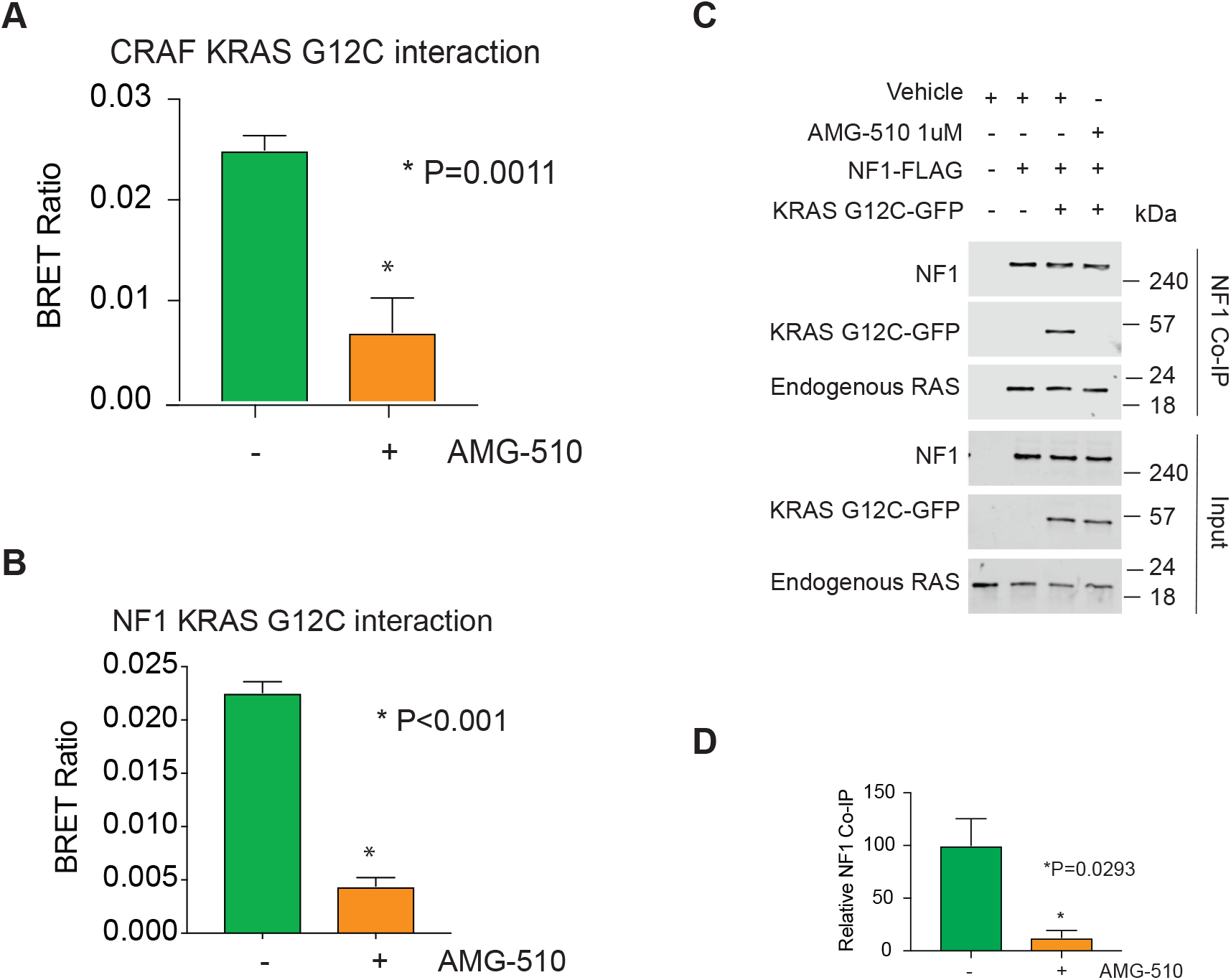
Experimental analysis of the effects of AMG-510 bound KRAS-G12C cancer cells. (**A**) HEK293T cells were co-transfected with KRAS-GFP and NF1-NanoLuc with and without 500nM AMG-510 for 24 hours and signal is represented as a BRET ratio for both sample groups. Bars represent the mean of eight biological replicates. Results are representative of an individual experiment from three separate experiments. (**B**) HEK293T cells were co-transfected with KRAS-GFP and CRAF-NanoLuc with and without 500nM AMG-510 for 24 hours and signal is represented as a BRET ratio for both sample groups. Bars represent the mean of eight biological replicates. Results are representative of an individual experiment from three separate experiments. (**C**). HEK293T cells were transfected with either KRAS-G12C-GFP and NF1, NF1 alone, or mock transfection. Cells were then treated with either vehicle or 500 nM AMG-510 for 24 hours. Following treatment, cells were lysed and NF1 co-IP was performed. IP product and input lysates were resolved and subjected to western blot analysis. Results are representative of an individual experiment from three separate experiments. (**D**) Mean KRAS G12C pull-down for the three independent NF1-coIP experiments. Statistical analyses were performed with unpaired t-test and P-values are indicated. Error bars in all panels represent standard deviation.

**Supplementary Figure 2.**
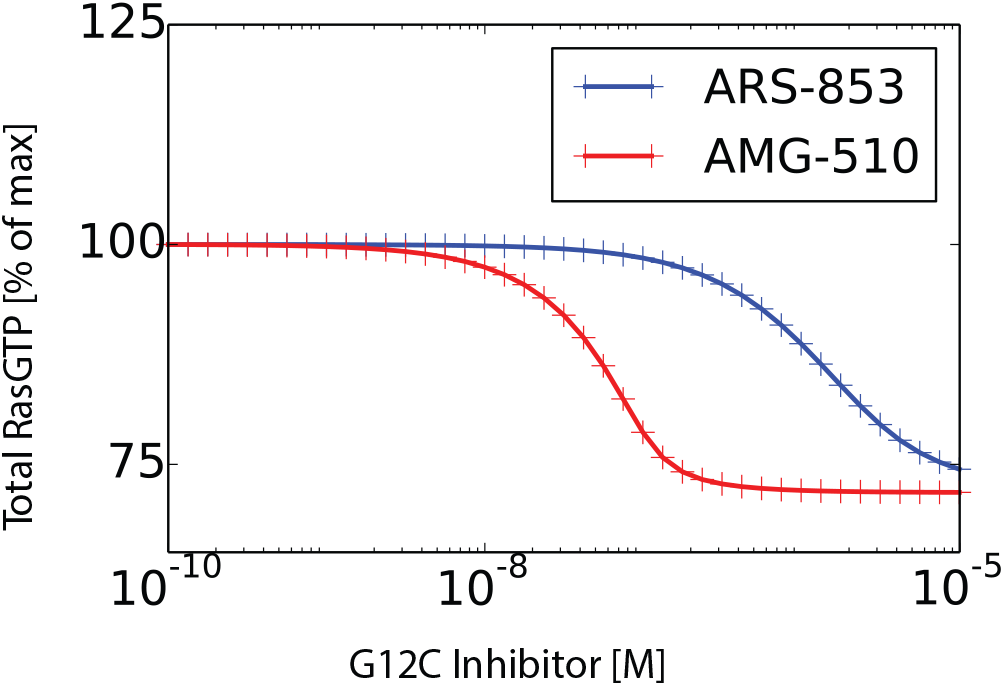
KRAS G12C inhibition mathematical model predictions for different KRAS G12C inhibitors. Simulated G12C inhibitor dose responses from the computational RAS model for KRAS G12C for AMG-510 and for ARS-853.

## Materials and Methods

### Cell line Models and Culture

SW837, SW1463, SW48 cells and SW48 isogenic counterparts were cultured in RPMI 1640 medium supplemented with fetal bovine serum (FBS) (10%), penicillin (100 U/ml), streptomycin (100 μg/ml), and l-glutamine (2 mM). All cells were grown in indicated medium and incubated at 37°C in 5% CO2 unless indicated otherwise in experimental methods. SW48 cells were obtained from Horizon Discovery. SW837 and SW1463 cells were obtained from the American Type Culture Collection.

### Proliferation Assay

Cells (5000 per well) were seeded in 96-well plates in complete media. Treatments were initiated after the cells were attached (24h). At the appropriate time points, cell viability was determined by MTT assay; (5 mg/ml in phosphate-buffered saline) was added to each well followed by incubation at 37°C for 2 hours. The formazan crystal sediments were dissolved in 100 μl of dimethyl sulfoxide, and absorbance was measured at 590 nm using a Tecan Infinite 200 PRO plate reader. Each treatment was performed in eight replicate wells and repeated three different times.

### Western Blot Analysis

Cell lysates were generated using lysis buffer (Thermo Fisher Scientific, 1862301) containing protease inhibitor cocktail (Cell Signaling Technology) and incubated on ice for 1 hour, with brief vortexing every 5 minutes. The total protein concentration was determined by Pierce Protein assay (Thermo Fisher Scientific). Protein samples were resolved by electrophoresis on 12% SDS–polyacrylamide gels and electrophoretically transferred to polyvinylidene difluoride (PVDF) membranes (Millipore Corporation) for 20 min at 25 V with the Trans-Blot Turbo (Bio-Rad Laboratories). The blots were probed with the appropriate primary antibody and the appropriate fluorophore-conjugated secondary antibody. The protein bands were visualized using the Licor CLx Odyssey imaging station (Licor Biosystems). Comparative changes were measured with Licor Image Studio software from three independent experiments. Comparisons were made by normalizing to endogenous loading control for internal reference, and to control treatment for external reference.

### AMG-510 NF1 Coimmunoprecipitation

HEK293T cells were individually transfected or co-transfected with the expression plasmid for NF1-Flag, WT KRAS-GFP, or G12C KRAS-GFP. Cells were also simultaneously treated with either vehicle or 500nM of AMG-510 for 24h. Cells were harvested in IP Lysis/Wash Buffer (0.025 M tris-HCl, 0.15 M NaCl, 0.001 M EDTA, 1% NP-40, and 5% glycerol; pH 7.4 and 1× protease inhibitor) 24 hours after transfection and treatment. Whole-cell lysates (500 μg) were precleared for 0.5 hours using Control Agarose Resin slurry (Thermo Fisher Scientific). Immunoprecipitation was performed by first incubating 800 μl of HEK293T NF1-Flag precleared lysate with 200 μl of either WT KRAS-GFP, G12V KRAS-GFP or G12C KRAS-GFP precleared cell lysate. The final steps of the coimmunoprecipitation were performed using the Pierce Immunoprecipitation Kit (Thermo Fisher Scientific) with immobilized anti-NF1 antibody (Santa Cruz Biotechnology, CA). A total of 500 μg of the cell lysate was added and incubated at room temperature under rotary agitation for 45 m. At the end of the incubation, the complexes were washed five times with lysis buffer. The western blot was probed with mouse monoclonal NF1 antibody (Santa Cruz Biotechnology) and mouse monoclonal RAS antibody (Thermo Fisher Scientific).

### Bioluminescence Resonance Energy Transfer (BRET) assay

Human embryonic kidney HEK293T cells were grown in DMEM/10% FBS without antibiotic treatment. Cells were seeded at 5 × 100 cells per well in a 96-well white opaque Perkin Elmer microplate. Twenty-four hours after seeding, cells were co-transfected with either a constant concentration of 0.1 μg of NF1-NanoLuc pcDNA expression plasmid or CRAF RBD-NanoLuc pcDNA expression plasmid with 0.2 μg of GFP-tagged KRAS (WT or Mutant) with 0.25 μl of Lipofectamine 2000 per well following the manufacturer’s protocol (Thermo Fisher Scientific). Twenty-four hours later, medium was aspirated from each well and 25 μl of Nano-Glo Live Cell Reagent was added to each well per the manufacturer’s protocol (Promega). Plates were placed on an orbital shaker for 1 min at 300 rpm. After incubation, the plate was read on the Tecan Infinite M200 PRO with LumiColor Dual Setting with an integration time of 1000 ms. BRET ratio was calculated from the dual emission readings. BRET ratio was plotted as a function of the RAS-GFP/NF1-NanoLuc plasmid ratio. BRET assays were repeated three times, each with eight biological replicates.

### Excess Over Bliss Score

Cell proliferation index was converted to fraction affected (fA);(1 – percent viable)=fA. The predicted value (C) was calculated for each dose where A corresponds to fraction affected for cetuximab and B corresponds to fraction affected for AMG-510: C = (A+B) – (A x B). Excess over Bliss (EOB) was calculated as: EOB= (fA_(A+B)_-C) x100, where fA_(A+B)_ corresponds the fraction affected of combination of same dose of both Cetuximab and AMG-510.

### Active RAS Pull-Down Assay

Isolation of active RAS-GTP was performed using the Active Ras Pull-Down and Detection Kit (Thermo Fisher Scientific) following the manufacturer’s protocol. RAS abundance was measured by Western blot. Western blot analysis of RBD pull-down lysates was performed with mouse anti-KRAS antibody (WH0003845M1, Sigma), rabbit anti-NRAS (ab167136, Abcam), rabbit anti-HRAS (18295, Proteintech), mouse anti–pan-RAS antibody (1862335, Thermo Fisher Scientific), and mouse anti-GAPDH (sc-4772, Santa Cruz Biotechnology). Input lysates were analyzed with mouse anti-pERK (675502, Biolegend) and rat anti-ERK (686902, Biolegend).

### Mathematical Model

The RAS model utilized here is an extension of our original RAS model ^16^ that was extended to include protein production and protein degradation ^17^. We developed this extended model to study situations where protein turnover must also be considered, such as to study the effects of KRAS G12C inhibitors. The modeled half-life for RAS proteins (24 hours) is consistent with recent measurements ^8^. We use this version of the model to study combined treatment with KRAS G12C inhibitors and EGFR inhibitors. For both models, we model KRAS G12C to have impaired intrinsic GTPase activity (k_GTPase_ = 72% of the value of k_GTPase_ for WT RAS) ^23^, GAP mediated activation (k_cat_ equal to zero, with the assumption that codon 12, 13, and 61 RAS mutations are insensitive to GAPs, but with the GAP bound KRAS G12C-GTP protein still capable of intrinsic GTPase activity), and with a lower affinity for binding to RAS effectors (modeled as slightly faster dissociation from effectors, k_d,effector_ = 120% of the value of _kd,effector_ for WT RAS) ^23^, with all of these values based on previously published, experimental studies as cited. Based upon our separate detection of impaired binding to NF1 ^15^, we also modeled NF1 binding with an impaired binding to KRAS G12C (K_m,GAP_ = 1000% of the value of K_m,GAP_ for WT RAS). EGFR inhibitor dose responses are simulated with the assumption that EGFR mediated activation of RAS is driven by the recruitment of GEFs like SOS1 and SOS2 to activated EGFR, and that EGFR inhibition results in a reduction of this GEF mediated activation of RAS-GTP, as we have done previously ^13^. KRAS G12C inhibition is modeled as done previously ^17^, but with new second order KRAS G12C inhibitor covalent binding parameters specific for AMG-510 ^1^. Model simulations are used to determine steady-state levels of total RAS-GTP (KRAS-GTP + NRAS-GTP + HRAS-GTP) as a measure of RAS pathway signal output. Our model is a set of Ordinary Differential Equations (ODEs) that we solve numerically using the Python package SciPy. Programs to reproduce all computational results are attached as supplementary code.

## Acknowledgments

We thank Tony Hunter and the members of the Stites and Hunter laboratories for helpful conversations and feedback. We thank Shumei Kato for providing cetuximab, Max Shokhirev for statistical consulting, and Jose Mendoza Lopez for assistance with experiments. This work was supported by NIH K22CA216318, NIH T32CA009370, NIH P30CA014195, NIH DP2AT011327, and the Joe and Dorothy Dorsett Brown Foundation. Author contributions: T.M and E.S. designed the experiments and study. T.M. performed the experiments with assistance from L.S.H. T.M. performed the statistical analysis. M.T. and E.S. performed computational analyses. T.M., M.T. and E.S. wrote the manuscript with input from the other authors. The authors declare that they have no competing interests.

